# PresRAT: A server for identification of bacterial small-RNA sequences and their targets with probable binding region

**DOI:** 10.1101/2020.04.03.024935

**Authors:** Krishna Kumar, Abhijit Chakraborty, Saikat Chakrabarti

## Abstract

Bacterial small-RNA (sRNA) sequences are functional RNAs, which play an important role in regulating the expression of a diverse class of genes. It is thus critical to identify such sRNA sequences and their probable mRNA targets. Here, we discuss new procedures to identify and characterize sRNA and their targets via the introduction of an integrated online platform named PresRAT (Prediction of sRNA and their Targets). PresRAT uses the primary and secondary structural attributes of sRNA sequences to predict sRNA from a given sequence or bacterial genome. PresRAT also finds probable target mRNAs of sRNA sequences from a given bacterial chromosome and further concentrates on identification of the probable sRNA-mRNA binding regions. Using PresRAT we have identified a total of 60,518 potential sRNA sequences from 54 bacterial genomes and 2447 potential targets from 13 bacterial genomes. We have also implemented a protocol to build and refine 3D models of sRNA and sRNA-mRNA duplex regions and generated 3D models of 50 known sRNAs and 81 sRNA-mRNA duplexes using this platform. Along with the server part, PresRAT also contains a database section, which enlists the predicted sRNA sequences, sRNA targets and their corresponding 3D models with structural dynamics information.

## Introduction

small-RNA (sRNA) based regulation of mRNA sequences play a central role in various bacterial gene expressions in response to environmental changes^1, 2^. These sRNA sequences are extensively studied in context with different bacterial regulatory processes such as quorum sensing^3^, the formation of biofilms^4^, expression of diverse virulence factors^5^, iron uptake^6^, etc. In prokaryotic system, these functional RNA regulators exist as small transcripts about 30-600 nucleotides long and modulate the translation and stability of mRNAs by binding with them^7^. Binding regions of these sRNA sequences with their cognate mRNAs are not fixed. Some sRNA sequences primarily bind at the ribosome binding site (RBS) of their target mRNA sequences^1^ while other sRNA sequences modulate translation of target genes by attaching at distant locations^1,7^ The antisense mechanisms exhibited by the sRNA sequences to modulate their targets are the key to understand the function of these RNA regulators^1,2,7^ The significance of these sRNA sequences and their targeted genes in bacteria has driven the research community to identify and more of these sRNA sequences and their targets with respect to various biological functions. Different experimental methods such as RNA-seq, tiling arrays, sRNA pulse expression, immunoprecipitation are available to detect sRNA sequences and their targets but they are tedious and expensive and the outputs of these experiments also need exhaustive filtering^8–10^. Hence, computational approaches can be helpful in augmenting the experimental methods^11^ in the detection of sRNA sequences and their targets.

Comparative genomics is a standard computational practice to find sRNA sequences within a bacterial genome^12^. QRNA^13^ and Intergenic Sequence Inspector^14^ programs implemented a comparative genomics approach to find sRNAs. Programs like RNAz^15^ and sRNApredict^16^ use the conserved RNA secondary structure information to calculate the thermodynamic energy for sRNA prediction. GLASSgo^17^ combined an iterative BLAST strategy with pairwise identity filtering and structure-based clustering to find sRNA homologs from scratch. Some of the programs also utilize the promoter and terminator sequence signal annotations for sRNA prediction. sRNAscanner^18^ applies a position weighted matrix of sRNA specific transcriptional signals and sRNAfinder^19^ utilizes a generalized probabilistic model to integrate primary sequence data, transcript expression data, microarray experiments and conserved RNA structure information to to predict sRNA genes. Diverse sets of methods also utilizes different machine learning approaches^20,21^, antisense RNA transcript expression profiling^22^ and motif-based discovery^23^ algorithms to predict sRNA sequences.

Identification of sRNA target genes are also important and represents a challenging task. Experimental procedures including genetic screens^24^, knockouts studies^25^, microarray analysis^26^ and other systematic proteomic investigations^27^ are implemented to identify sRNA target genes. However, these methods are expensive, time-consuming and extremely tedious. Therefore, development of sensitive and accurate computational approaches would aid and complement the experimental analysis. The first step in sRNA target prediction is to search for a complementary region between the sRNA and its target mRNA sequence. Comparing to microRNA and its target mRNA interaction, which requires a generally conserved 6-8 nucleotide long seed region for initiation of binding, the sRNA-mRNA binding region is relatively more extended and more evolutionary flexible^28^. Non-Watson-Crick basepairing plays a key role in sRNA-mRNA target interaction and programs such as TargetRNA^26^, GUUGle^29^ utilises both Watson-Crick and non-Watson-Crick base pairing models to find sRNA targets and their interacting regions. RNAcofold^30^ computes the hybridization energy and base-pairing arrangement between a pair of RNA molecules, while RNAplex^31^ calculates the same hybridization energy using modified thermodynamic energy models. RNAup^32^ utilises the thermodynamics of RNA-RNA interactions and the hybridization energy while IntaRNA^33^ incorporates binding site accessibility information for searching the best interaction with minimum extended hybridization energy. SPOT^34^ incorporated existing computational tools to search for sRNA binding sites and utilises experimental data for filtering.

Despite the availability of various sRNA and its target prediction algorithms, the sensitivity remains quite low (20% – 49%)^11^. Hence, it is fair to assume that many more sRNA sequences and their target genes remain undiscovered, particularly in less studied prokaryotic organisms. Here, in this article, we introduce an integrated platform with alternative approach to identify bacterial small-RNA and their target genes from a given nucleotide sequence and from a whole bacterial chromosome. This online platform, PresRAT (<u>Pre</u>diction of <u>sR</u>N<u>A</u> and their <u>T</u>argets) integrates two separate but related procedures of sRNA and their target identification into a simple and effective single dais.

Apart from identifying the sRNA sequences and their targets, a key feature of this platform is to provide sRNA and sRNA-mRNA duplex structures generated via three dimensional (3D) modelling and further refined through molecular dynamics simulation. Predicting accurate 3D structure of RNA from the primary sequence is still a fundamental and challenging problem in computational biology^35^. There are few notable programs such as iFoldRNA^36^, FARNA/FARFAR^37, 38^, MC-Sym^39^, MANIP^40^, 3DRNA^41^, RNA2D3D^42^, NAST^43^ and ASSEMBLE^44^ try to tackle the existing bottlenecks of RNA structure prediction and improve the algorithms for RNA 3D structure generation. As per the latest RCSB protein databank^45^ statistics, there are 1528 experimentally solved 3D structures of RNA representing only ~0.01% of the whole deposited structures. So far there is only one experimentally solved 3D structures available for sRNA^46^ (N-terminal SL1 part of DsrA from *E.coli)* other than few sRNA conjugated with chaperone proteins^47–51^. PresRAT uses RNA2D3D^42^ program to generate an initial RNA model followed by extensive refinement of the structure using GROMACS v4.6.3 molecular dynamics simulation package^52^.

The validation and benchmarking results of the PresRAT for sRNA, sRNA-target, and sRNA-mRNA binding regions suggest improvements compared to those achieved by other available programs. In this current study we have studied 54 bacterial genomes to identify a total of 15,568 and 44,950 sRNA sequences from genic and non-genic regions of the respective genomes. We also have identified 2447 potential targets for 50 unique sRNA sequences from 13 bacterial genomes. Further, we have shown the structural details of 50 sRNA and 81 sRNA-mRNA duplex 3D models through molecular dynamics simulation studies. All these procedures and information are automated and integrated in the PresRAT platform freely available at **http://www.hpppi.iicb.res.in/presrat**.

## Results

### PresRAT: The webserver interfaces

PresRAT webserver and the database sections are built using CGI-Perl scripts. Integrated CGI-Perl scripts allow the users to utilize this webserver in an efficient way. The webserver is freely available at www.hpppi.iicb.res.in/presrat. The PresRAT webserver section is divided into four parts accomplishing their individual tasks. The first two parts are called **Pred^sRNA^** and **Pred^GsRNA^**. **Pred^sRNA^** interface can be used to identify sRNA sequences one at a time while **Pred^GsRNA^** interface directs the user to identify potential sRNA sequences from different bacterial sequences. The next two parts of the PresRAT webserver compromises sRNA target identification protocols. **Pred^Tar^** Interface helps to identify interacting nucleotides between a pair of sRNA and mRNA sequence and **Pred^WG^** performs the function of sRNA target identification from a given bacterial genome. The following sections briefly describe the methodology and the results of each of these parts of the PresRAT server.

#### Pred^sRNA^: Prediction of sRNA from its sequence and secondary structural attributes

**Pred^sRNA^** uses sequence and secondary structural information of existing sRNA and non-sRNA sequences to calculate a combined score to predict novel sRNA sequences. In sequence analysis, a directional (5’->3’) dinucleotide score is calculated for the whole input sequence from Log Odds (LOD) score matrices of all possible combinations of directional dinucleotides, which we denote as *Sequence_score_* (see sRNA prediction protocol and Figure S1 in Supplementary Information File). We also calculate a *U-rich_score_* that reflects the presence of uracil (U) nucleotides at the 3’ end of the sRNA sequence. PresRAT effectivelyuses the secondary structure information by first calculating a set of local minima conformations of the provided RNA sequenc. followed by estimating the two different scores namely *ABE_score_* and *ALE_score_. ABE_score_* and *ALE_score_* indicate the Average Base Energy and Average Loop Energy of local minima secondary structure conformations. The final combined score i.e. *sRNAscore* is calculated as follows:

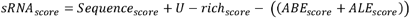

The combined *sRNAscore* (details of the scoring schemes can be found in Supplementary Information File) is used as the final scoring scheme for filtering the true positive sRNA instances from the true negative ones.

**Pred^sRNA^** page accepts query nucleotide sequence and based on its *sRNA_score_* and predicts whether the given sequence is likely to be sRNA or not above a user defined confidence level. The output page provides the *sRNAscore* and its other component’s score along with a distribution of the score with respect to the known sRNA scores. **Pred^sRNA^** result page also provides a minimum free energy secondary structure of the probable sRNA sequence. **Pred^sRNA^** also provides a useful option of building the threedimensional (3D) structural model of the predicted sRNA sequence via the **Pred^sRNA3D^** module, which generates sRNA model using RNA2D3D^42^ and RNAfold^53^ programs, respectively. Following the generation of a 3D sRNA structure, a molecular dynamics (MD) simulation is performed on the RNA structure using GROMACS^52^ MD simulation package (details of the structural analysis can be found at Supplementary Information File). The initial and final structures are provided from MD simulation trajectory file along with other necessary information.

For benchmarking and validation of sRNA prediction by PresRAT, a set of non-redundant 819 sRNA (Additional file 1), 31,403 genic, and 28,733 non-genic non-sRNA sequences were randomly collected from 54 different bacterial species (Supplementary Information File and Additional file 1). Receiver operating characteristic (ROC) statistics were calculated by comparing the sRNA to non-sRNA *sRNA_score_* in a 1:5 ratio via 1000 fold crossvalidation protocol. In each cross-validation step sRNA and non-sRNA sequences were randomly collected to calculate the sensitivity and error rates. The ROC statistics generated using the *sRNA_score_* distribution of sRNA and non-sRNA sequences demonstrate sensitivity values of 13.2%, 28.0%, 39.0% and 50.1% at 1.0% 5.0%, 10.0%, and 20.0% error rates, against non-sRNA sequences collected from genic regions whereas sensitivities of 6.3%, 18.4%, 31.2%, and 48.0% were obtained at 1.0% 5.0%, 10.0%, and 20.0% error rates against non-sRNA sequences collected from non-genic parts of the bacterial chromosomes (Figure 1A), respectively. A comparative study against RNAz^15^ program with PresRAT showed slightly better or comparable performance in predicting sRNA sequences with respect to genic non-sRNA sequences (Figure 1B) while PresRAT showed much better performance in sRNA prediction with respect to non-genic non-sRNA sequences (Figure 1C). However, it is also important to mention that RNAz requires evolutionary information from homologous sequences to predict functional RNA sequences while PresRAT does not require homologous sequences for sRNA prediction. However, to make comparison meaningful between the two methods, comparative analysis was performed on a limited set of commonly predicted sRNA (199 sequences), genic (5193) and non-genic (4218) non-sRNA sequences as RNAz^15^ was restricted only to the mentioned number of sRNA and non-sRNA sequences having homologous sequences in same or other bacterial chromosomes.

**Figure 1.**
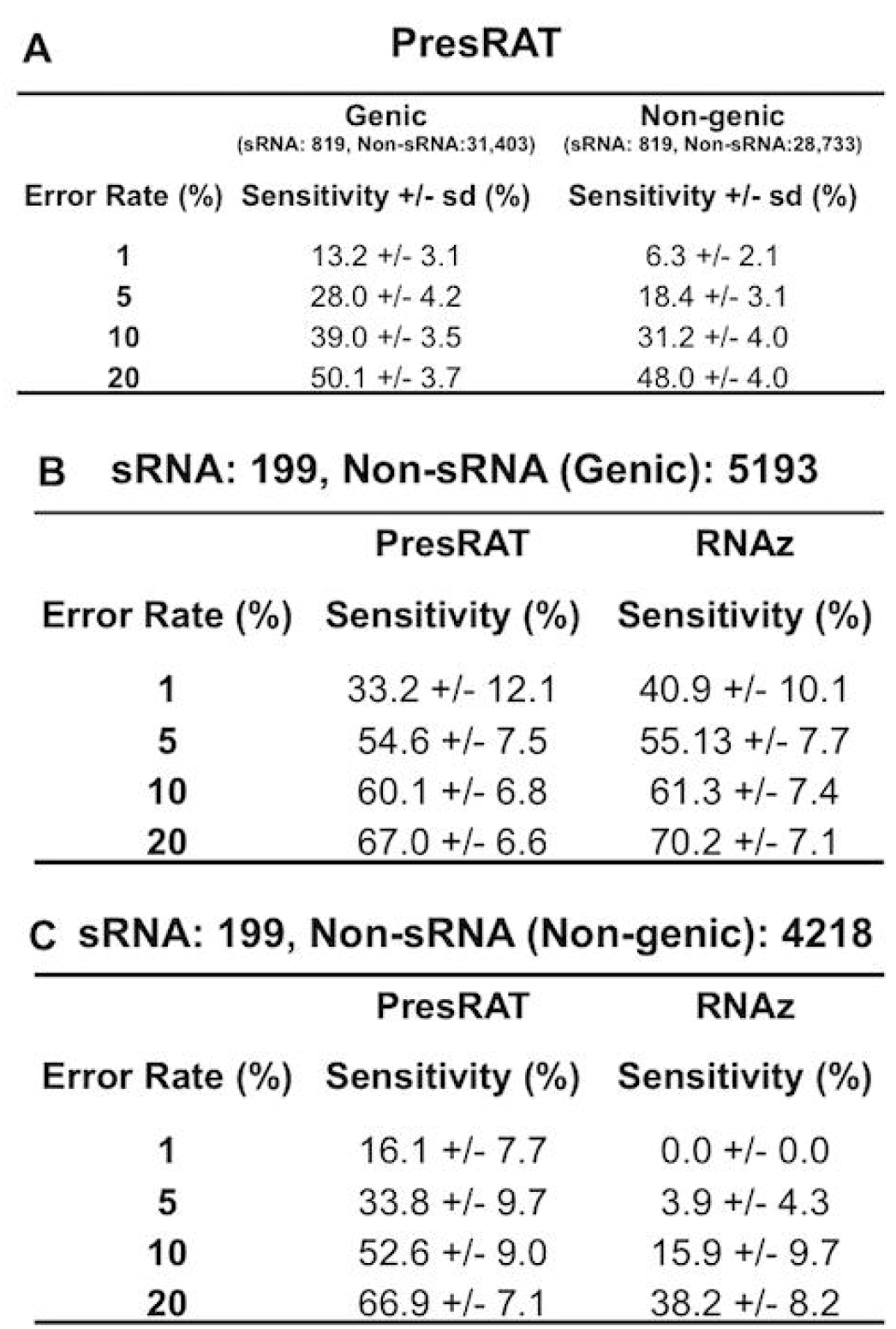
Panel A provides the ROC statistics of PresRAT program on the full dataset. Panel B and C show the ROC statistics perform on the limited set of commonly predicted sRNA and non-sRNA sequences.

**Pred^GsRNA^:** Prediction of sRNA sequences within bacterial chromosome **Pred^GsRNA^** aims to predict potential small-RNA (sRNA) sequences within a bacterial genome. The result page of **Pred^GsRNA^** contains the sRNA sequences predicted from the genic sequences and non-genic sequences of the bacterial genome, respectively. The corresponding result page of this program consists of top 100 predicted sRNA sequences along with their start and end loci, *sRNA_score_* and P-values. Similarly, a pictorial representation of the predicted sRNA sequences on a genome browser view is also provided for better visualization (Please refer the supplementary material for further details of this approach).

**Pred^GsRNA^** was used to predict new probable sRNA sequences from the 54 bacterial genomes (Additional file 2) to identify a total of 15,568 and 44,950 hypothetical sRNA sequences from genic and non-genic parts of the bacterial genomes (Additional file 3). sRNAscanner^18^, a separate program for bacterial chromosomal sRNA detection was used for comparative analysis. sRNAscanner uses a transcriptional signal based computational method for sRNA detection and detected only 55 unique sRNA sequences while our new protocol identified a total of 205 sRNA sequences out of 819 sequences (Additional file 4). Our protocol demonstrates moderate improvement and provides an alternative approach in identifying sRNA sequences from bacterial genomes.

#### Pred^TAR^: Prediction of sRNA-mRNA target binding regions

**Pred^TAR^** aims to predict potential binding regions between bacterial small-RNA (sRNA) and the target mRNA of that sRNA. The output page provides the alignment of the given sRNA and mRNA sequences along with scores and duplex energy values. PresRAT uses combines two different approaches to find a sRNA-mRNA target binding region. The first approach (Component I) uses the base un-pairing probability of both the sRNA and its target mRAN sequence (Figure 2A) and the subsequent approach (Component II) (Figure 2B) utilizes the local minima energy structures of sRNA and the minimum free energy structure of the target mRNA to find the potential binding region (Figure S2 in Supplementary Information File for details). **Pred^TAR^** also provides a useful option of building the three-dimensional structural (3D) model of the sRNA-mRNA bound duplex region via the **Pred^dup3D^** module, which generates the duplex model using RNA2D3D^42^ program for the secondary structure information followed by molecular dynamics (MD) simulation. Details of the structural analysis can be found at Supplementary Information File. The Initial and final structures are provided from MD simulation trajectory file along with other necessary information.

**Figure 2.**
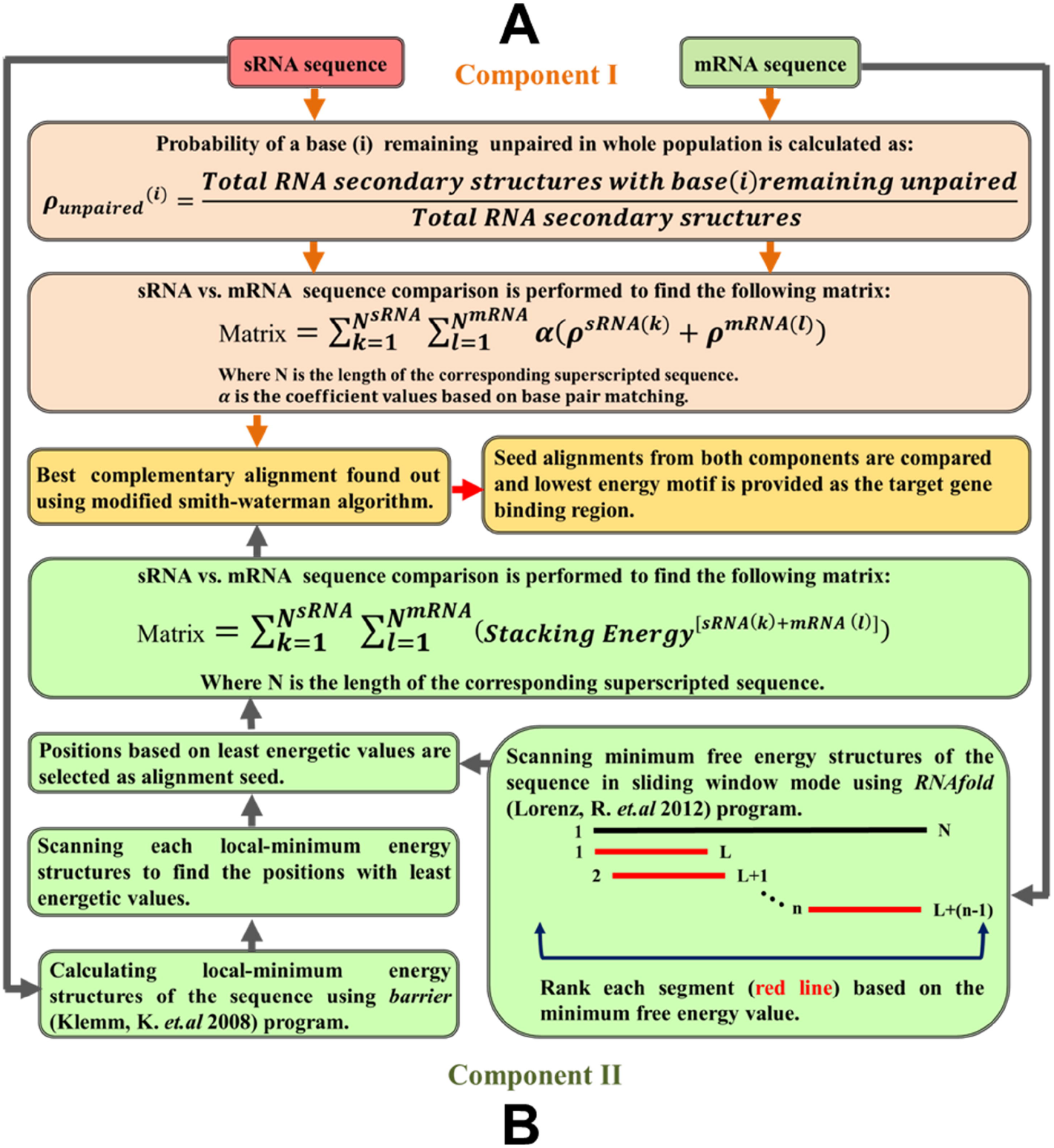
Panel A demonstrates the sRNA-mRNA target binding region identification protocol (Component I) of PresRAT. The main aim of this protocol is to identify the un-pairing probability of each base from both sRNA and mRNA sequences and utilise this probability information to identify the binding region. Panel B describes the Component II protocol used by PresRAT to identify sRNA-mRNA target binding region. In this protocol energetic value of both sRNA and mRNA secondary structures are taken into account to identify binding regions.

Experimentally known binding regions in 88 out of 91 sRNA and their respective target mRNA sequences (Additional file 5 and Additional file 6) were used to validate the Component I and II algorithms. The selection of these 88 binding regions was done based on the frequency distribution of the sRNA binding site on mRNA sequences relative to its translation initiation site (Figure 3A). The frequency distribution of the 91 sRNA binding sites on mRNA sequences reveals that in 97% of the cases the sRNA binding site is located within the −250 to +100 window region of a target mRNA (“0” being the translation start site). Component, I and II algorithms, could correctly identify the binding region for 49% and 67% of the true sRNA-mRNA pair, respectively within the above mentioned window region. The Figure 3B shows a Venn diagram illustrating the distribution of correctly predicted sRNA-target binding region between the two algorithms Component I and II. 74% (65 out of 88) sensitivity was achieved when the binding region with lowest free energy of sRNA-mRNA binding^54, 55^ predicted by either of the algorithms was considered. 82% (72 out of 88) sensitivity (Figure 3C) was achieved when both the common and uniquely predicted binding regions were considered. A comparative analysis (Figure 3C) with other published methods shows much better performance of PresRAT algorithms in predicting sRNA-mRNA targets (a complete list of predicted sRNA-mRNA binding regions is provided in Additional file 7).

**Figure 3.**
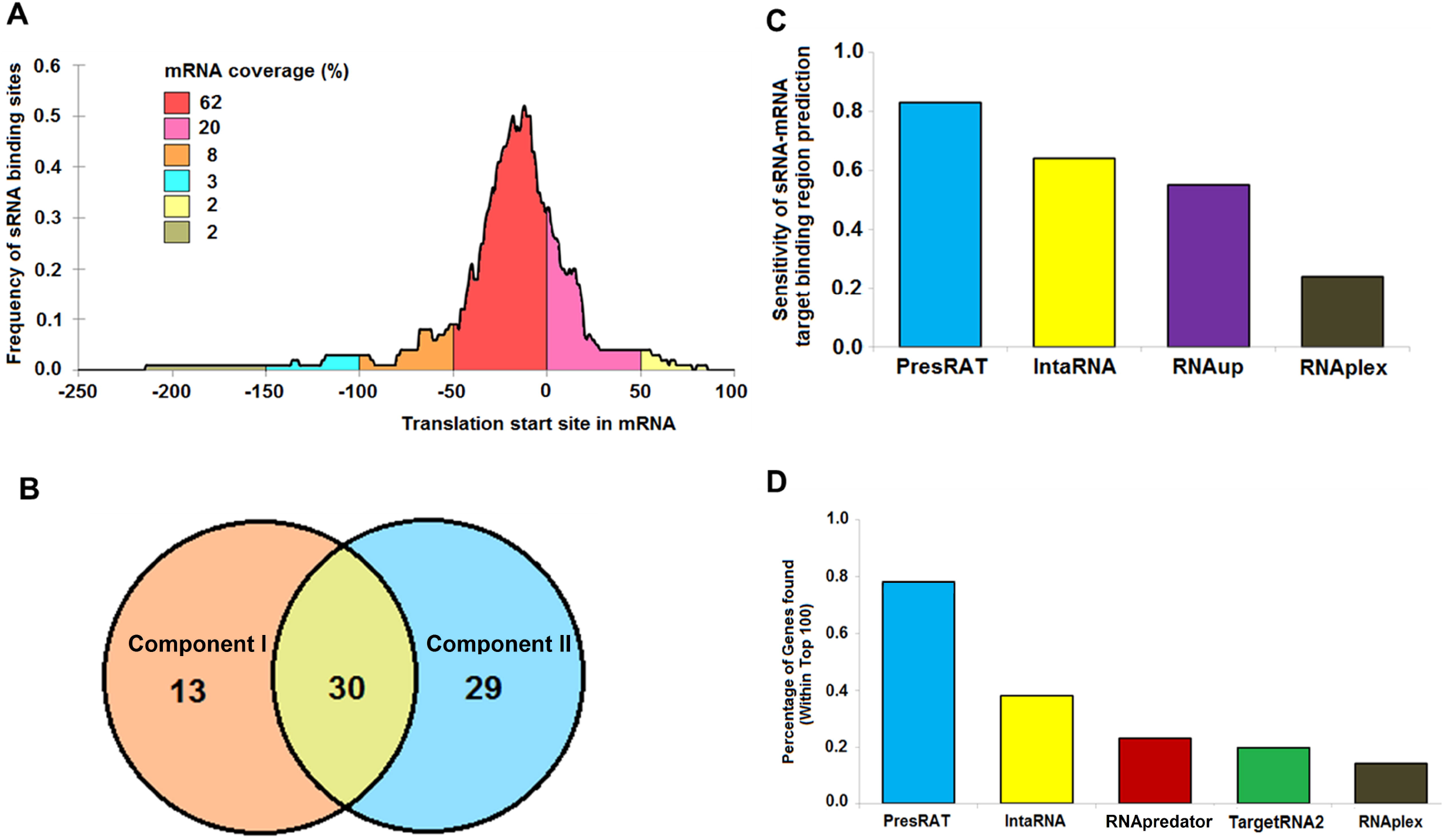
Panel A shows the frequency distribution of sRNA binding sites in the 91 mRNA sequences. In 98% of the cases it is found that the sRNA binding site is present within the window of −250 to +100 nucleotides (0 being the translation initiation site) of the target gene. Panel B shows the overlap of correctly predicted target by the PresRAT Component I and II protocols, respectively. Panel C shows the PresRAT sensitivity in predicting the correct sRNA-mRNA target binding regions in 88 samples compared to other programs. Panel D shows the percentage of true sRNA target gene identification through whole-genome search by PresRAT compared to other existing programs.

#### Pred^WG^: Prediction of sRNA target genes and binding regions within a bacterial genome

**Pred^WG^** aims to predict potential bacterial small-RNA (sRNA) targets within a bacterial genome. The result page of **Pred^WG^** contains the list of the predicted target genes along with individual alignment and the corresponding duplex energy value of the binding region. Similarly, a pictorial representation of the predicted sRNA sequences on a genome browser view is also provided for better visualization.

Our benchmarking exercise considering 91 known sRNA-mRNA target pairs, PresRAT achieved 79% sensitivity in rightly predicting the target genes (within the top 100 predictions/genome) within bacterial genomes compared to similar programs, like IntaRNA^33^, RNAPredator^56^, TargetRNA2^57^ and RNAplex^31^, which achieved sensitivity values of 38%, 23%, 22% and 14%, respectively (Figure 3D) (complete list of targets found by the various programs are provided in Additional files 6 and 7). For a fair comparison, a similar search window (−250 to +100 of mRNA translation start site) length was maintained for all the programs. Within the top 100 predictions for individual sRNA sequences, we have predicted 2447 novel mRNA targets with a similar binding energy distribution profile *(p* values < 0.5) (Figure S3 in Supplementary Information File) as that of the known 91 sRNA-mRNA interactions. The complete list of the identified target mRNAs for 50 sRNA sequences along with their corresponding functions is provided in the Additional file 8. Gene Ontology (GO)^59^ based molecular functional network of E. coli known sRNA sequences (18 sequences) with their known (27 unique targets) and predicted mRNA target sequences (192) (Figure S4A in Supplementary Information File) suggests involvement of similar functions such as binding, catalysis and transporter activities (Figure S4B in Supplementary Information File). Additionally, a smaller subset of the predicted target genes also offers unique molecular functions like electron transport, guanyl-nucleotide exchange and other regulator activities (Figure S4B in Supplementary Information File).

### PresRAT: The database interface

PresRAT database section enlists the predicted sRNA sequences, sRNA targets, and the corresponding 3D models of selected sRNAs and sRNA-mRNA duplex structures. Predicted small-RNA sequences from 54 bacterial genomes are accessible to users from a drop-down selection menu provided in the database page. Similarly, the list of predicted target genes for a given sRNA within a bacterial genome is readily accessible in a similar manner. The database also harbors mRNA targets of 50 known sRNA sequences from 13 bacterial genomes along with their details.

Exhaustive structural analysis was performed to predict the 3D structures of the known sRNA sequences and their corresponding binding regions with the target genes. Three-dimensional modelling of sRNA sequences and the target binding regions using their current secondary structure information provides the opportunity to gain structural insight, which in turn may become quite valuable in RNA based therapeutics development. There are 50 sRNA models in PresRAT database. All the sRNA and 81 unique sRNA-mRNA bound duplex models are explicitly refined using GROMACS package through one-nanosecond of molecular simulation runs. The final models with other structural detailsare available to users in the database section. Further methodical details and validation results can be found in Supplementary Information File.

## Discussion

Here, we introduce a web-based platform PresRAT to identify and analyze bacterial small-RNAs using both individual and genomic sequence level searches. The platform integrates a combination of scoring schemes developed based on primary and secondary structural attributes of small-RNAs. Using PresRAT we have scanned a total of 54 bacterial genomes to identify 15,568 and 44,950 potential sRNA sequences (Supplemental material 3). One of the key features of this platform is the generation of sRNA threedimensional models and their dynamic analysis. For in-depth structural analysis, availability of energy minimized and refined three-dimensional models of sRNAs are essential. PresRAT provides an option for automated model building and molecular dynamics simulation of sRNA in presence of water. Using this automated procedure we have successfully built 50 different small-RNA structures and analyzed some of their basic properties calculated from their dynamics data. Comparison of our server derived model of DsrA and the NMR structure of the SL1 of the DsrA (pdb code: 5WQ1)^46^ shows high similarities (RMSD: 2.75Å) between the structures, indicating reliability of our 3D model (Figure S7 in Supplementary Information File). However, the NMR structure covers only a short fraction of the DsrA sequence.

PresRAT enables a user to search for probable target mRNAs of sRNA sequences from a given bacterial chromosome and further concentrate on identification of the probable sRNA-mRNA binding regions. For identification of target binding regions, PresRAT uses base un-pairing probability information of sRNA-mRNA sequences and local minima of sRNA secondary structures. Both the approaches combined with other additional steps provide a simple but effective way of identification of target mRNA sequences from the whole bacterial chromosomes. In this way we have scanned 13 bacterial genomes and identified 2447 novel mRNA target sequences for an existing 50 sRNA sequences (Additional file 7). The identified target mRNAs for a given sRNA are provided in the database section of PresRAT along with their interacting nucleotide region and their related functional information. Existing model generation and molecular dynamics simulation procedure are extended to build the energy minimized and refined three-dimensional models of the sRNA-mRNA duplex regions, as well their structural characterization were also performed. Our study revealed that in general these duplex regions are thermodynamically unfavourable, which indicates the involvement of other critical factors for generation of stable sRNA-mRNA complexes.

## Acknowledgement

SC acknowledges CSIR-IICB for infrastructural support, CSIR Network Project (BSC-0121), Systems Medicine Cluster (SyMeC) grant (GAP357) from Department of Biotechnology (DBT), High Risk High Reward Grant (HRR/2016/000093) from Department of Science and Technology (DST) for financial supports.

## Additional Information

### Author Contributions

K.K worked on the web server and drafted the manuscript. A.C. collected and organized the data, developed the program, webserver and drafted the manuscript. S. C. drafted the manuscript and coordinated the project. All authors read and approved the final manuscript.

### Competing financial interests

The author(s) declare no competing financial interests.

