## Supplementary Information File for "PresRAT: A server for identification of bacterial small-RNA sequences and their targets with probable binding region"

\# Equal authorship.

*Corresponding author.

Running title: Identification of bacterial sRNA and their targets.

Keyword: sRNA identification; sRNA target prediction; RNA structure; RNA-protein interaction; RNA-RNA interaction.

**Dr. Saikat Chakrabarti**

Structural Biology and Bio-informatics Division

CSIR-Indian Institute of Chemical Biology

Contact:

**sRNA prediction**

***sRNA sequence collection***

In this study we have used 819 non-redundant known small-RNA (sRNA) sequences collected from 54 bacterial chromosomes. All these experimentally verified sRNA sequences were collected from sRNATarBase^1^ and Bacterial Small regulatory RNA database^2^. The distribution of these 819 sRNA sequences across different bacterial species is provided in the Additional file 1.

***Non-sRNA sequence collection***

The non-sRNA sequences were also collected from the same 54 bacterial chromosomes and were primarily divided into two different categories (models), non-sRNA sequences from genic region and non-genic regions. A total of 31,403 genic and 28,733 non-genic non-redundant, non-sRNA sequences were randomly collected from the 54 bacterial chromosomes mentioned in Additional file 3. Genic and non-genic region information for the corresponding bacterial chromosomes was obtained from the NCBI RefSeq database^3^. Repeat sequences and known sRNA sequences from these 54 bacterial chromosomes were not included in the non-sRNA dataset. Length distributions of non-sRNA sequences were also maintained similar to the known sRNA sequences.

***sRNA prediction protocol***

The protocol for sRNA prediction is based on combinations of four different scoring schemes, which are described below.

a. ***Sequence score calculation***

At first, the number of a nucleotide (*e.g.,* A) followed by another nucleotide (*e.g*., U) is calculated using the following formula

| **5` to 3` direction** | **sRNA** | **Non-sRNA** |
| --- | --- | --- |
| **A followed by U** | **a** | **b** |
| **A Not followed by U** | **c** | **d** |

Followed by a log odds ratio is calculated by

$Log Odds Ratio=log\left/ \frac{a\times d}{b\times c} \right.$	(1)

All the individual directional (5' to 3') dinucleotide Log Odds Ratio are then calculated for both the sRNA (+) and non-sRNA (-) model datasets using the above formula (1).

A positive log odds ratio score indicates a higher chance of observing a specific dinucleotide combination and *vice-versa*. Likewise Log Odds Ratio scores of all the 16 specific dinucleotide combination are calculated against both genic and non-genic non-sRNA sequences (Figure S1). The final *Sequence_score_* is the sum of all directional dinucleotide log ratio scores calculated from the 16 nucleotide matrix divided by the length of the sequence.

${Sequence}_{score}= \frac{\sum_{i}^{L-1} {\overset{\to}{{(N}_{i}N_{i+1})}}_{score}}{Length of the sequence (L)}$	(2)

Where,	*L* is the length of the sequence, *N_i_* is the first base and *N_i+1_* is the second base of the sequence in 5' to 3' direction.

**Figure S1**

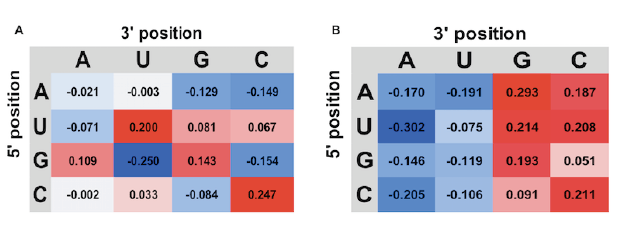

**Figure S1.** Panel A and B shows the Log-Odds ratio calculated for dinucleotide combinations of sRNA sequences against genic and non-genic sequences. A positive score (red color) presents a preferred dinucleotide in the sRNA compared to genic or non-genic sequences while a negative score (blue color) represents a non-preferred dinucleotide combination in sRNA sequences.

***b. Average base energy score calculation***

Energy landscape of a biomolecule provides insights towards the thermodynamic cooperativity during its folding pathway^4^. The shape of this landscape can illustrate the presence of stable intermediates (at minima) or barriers dividing the intermediates (at maxima). From previous studies^4, 5^ it is known that RNA molecules exhibit functional folding intermediates. Using this background knowledge, we explored the intermediate secondary structures of sRNA sequences to derive useful information required for their identification. The energy landscape of a RNA molecule is a two dimensional surface defined by the free energy versus conformational adaptations of the molecule^4, 6, 7^. A local minima RNA structure^8^ within this energy landscape is described as a secondary structure having energy lower than the energy of all neighbouring secondary structures. The decomposition of this energy landscape into local minima structures within a basin connected by saddle points generates the macro-states of the RNA molecules^9^. The macro-states or local minima structures are used to calculate the *Average Base Energy (ABE) score* of a RNA sequence. An *ABE_score_* is the average free energy values of all the local minima secondary structures divided by the length of the RNA sequence. The *ABE_score_* indicates the average contribution of each base in forming the whole local minima population of RNA secondary structures within the energy landscape.

***c. Average loop energy score calculation***

Loops generally provide thermodynamic stability to the overall structure of the RNA^10^. Here we have calculated the *Average Loop Energy (ALE) score* from a population of local minima secondary structures of a RNA molecule. *ALE_score_* suggest the stability of the RNA loops within a local minima secondary structural population. Lower the *ALE_score_* value, more the stable the RNA loops are. *ALE_score_* is calculated in the following way

${ALE}_{score}= \frac{1}{N} \sum_{i=1}^{n} \frac{{Total loop energy of structure}_{i}}{{Total number of loop in structure}_{i}}$	(3)

Where, N is the total number of local minima secondary structures in the population and *i* is the number of individual RNA secondary structures present within the local minima population.

***d. Terminal U rich score calculation***

A terminal “U” nucleotide (last 10 bases of each RNA sequences) richness score is calculated by the following formula

${U-rich}_{score}=\frac{\frac{Total U count\left( last 10 bases \right)}{10}}{\frac{Total U count}{Length of the RNA sequence}}$	(4)

The final score for sRNA to non-sRNA discrimination is calculated in the following way:

$\begin{aligned} \\ {sRNA}_{score}={Sequence}_{score}+{U-rich}_{score}- \left( \left( {ABE}_{score}+{ALE}_{score} \right) \right) \end{aligned}$	(5)

A positive and higher *sRNA_score_* can distinguish a sRNA from a population of non-sRNA sequences.

**sRNA-mRNA target site prediction**

***sRNA-mRNA target pair data collection***

91 unique pairs of sRNA-mRNA target pairs having the information of exact binding region at the nucleotide level were collected from 13 bacterial chromosomes to study and validate the target site prediction module. Additional file 5 lists those 91 sRNA-mRNA target pairs respectively. All the pairs are experimentally validated and acquired from sRNATarBase database^1^.

***Prediction of sRNA-mRNA target binding region***

We have used two different but complimenting approaches to find the sRNA-mRNA target binding region. The first approach uses the base un-pairing probability information of both the sRNA and its corresponding target mRNA (Component I). The Other approach (Component II) utilized the local minimum energy of sRNA secondary structures and the minimum free energy structure of the target mRNA to find the potential binding region. Both the components employed Smith-Waterman algorithm^11^ to find out the best binding region between the sRNA and its target mRNA. Descriptions for individual algorithms are provided below.

a. ***Predicting sRNA-mRNA binding using base un-pairing probability (Component I)***

In general, in order to get hybridized intermolecular base-pairing between two RNA molecules requires two un-paired regions^12^ within their respective sequences. So, to locate the probable RNA-RNA hybridization positions, detection of such un-paired regions within the RNA secondary structures is necessary. In this algorithm we first calculated the base un-pairing probability of both sRNA and mRNA sequences by first estimating the partition function of RNA sequences within a restricted base-pairing span^13, 14^ and then computing the mean probability of region of length 1 (one) remaining unpaired within a window span of sequence in the RNA suboptimal secondary structures^13^. This provides a list of un-pairing probability values for individual bases from both sRNA and mRNA. A sequence comparison between the sRNA and mRNA is performed using the modified Smith-Waterman algorithm to obtain a stretch of potential intermolecular complementary base-pairs. The potential score of a pair between a sRNA and mRNA base is the sum of probability values of the respective bases. In final Component I score calculation the above score is multiplied with a base-pair match-mismatch value. In canonical (A-U & G-C) and non-canonical (G-U) base-pair this value is set to 1 and 0.5, respectively, whereas for mismatched base-pair it is set to be -1. Therefore, a stretch of intermolecular base-paired region between sRNA and mRNA represents the best possible hybridized region having the highest base un-pairing probability of the respective RNA secondary structure population.

b. ***Predicting sRNA-mRNA binding using RNA local minima secondary structures (Component II)***

A local minima RNA structure^8^ is defined as a secondary structure having energy lower than the energies of all neighbouring secondary structures in its energy vs. conformational landscape. The total number of local minima secondary structures of a RNA sequence is constrained by the length and nature of the bases present in the sequence. Thus, instead of searching the whole suboptimal structures for un-paired region, in Component II algorithm we restrict the search within a pool of local minima secondary structures to find the least energetically stable structural region. The local minima secondary structures of sRNA are calculated using barrier program^8^, RNAfold^13, 15^ program is used to scan the mRNA sequence in a window basis (70 nucleotides) to rank and find the highest minimum free energy structure. A pool of sRNA local minima secondary structures and selected highest minimum free energy structure of mRNA are then individually scanned by the RNAeval program^13^ to select the least energetically stable region within their respective sequences. The pair with least energetically stable regions from both sRNA and mRNA is then compared by Smith-Waterman algorithm to find complementary base pairing. Unlike Component I, the Component II method uses only base-stacking energy values^16^ to obtain the most potentially stable stretch of intermolecular base-paired region between the sRNA and mRNA partners.

***Searching of sRNA target in whole bacterial chromosome***

The workflow developed to predict sRNA target genes in bacterial genome consists of number of steps. In the first step, a stretch of sRNA seed sequence is identified using both Component I and II algorithm. In case of Component I , the sRNA seed sequence represents the region having the highest un-pairing probability while for Component II the sRNA seed sequence reflects the least energetically stable region calculated from its local minima secondary structures. Next a complementary base-pair matching is done within the upstream (-250 to +100) sequences of the query genes using blastn program^17^. Once a seed sequence match is found against the query genes, the upstream sequences are searched against the known sRNA targets (upstream -250 to +100 nucleotide sequences). If satisfied with the filtering criteria for homology based searching (blastn score >= 30 and query coverage >= 30%) the upstream query gene sequences are forwarded to final score, potential binding region and ranking calculation by Component I and II algorithms. An alternative step is also applied to find the mRNA targets if all the above steps fail to find any suitable complementary region between sRNA and mRNA. In this alternative step potential mRNA seeds were derived from their corresponding DNA curvature profiles, followed by complementary base-pair matching with the sRNA sequence using blastn scan^17^. Once a match is found, a similar homology based approach is applied to select the target. Final target binding region between sRNA and mRNA is calculated using Component I and Component II method, mentioned previously. The final ranking of the target genes are done based on Component I and Component II scores. Figure S2 depicts the overall strategy of target identification procedure.

**Figure S2**

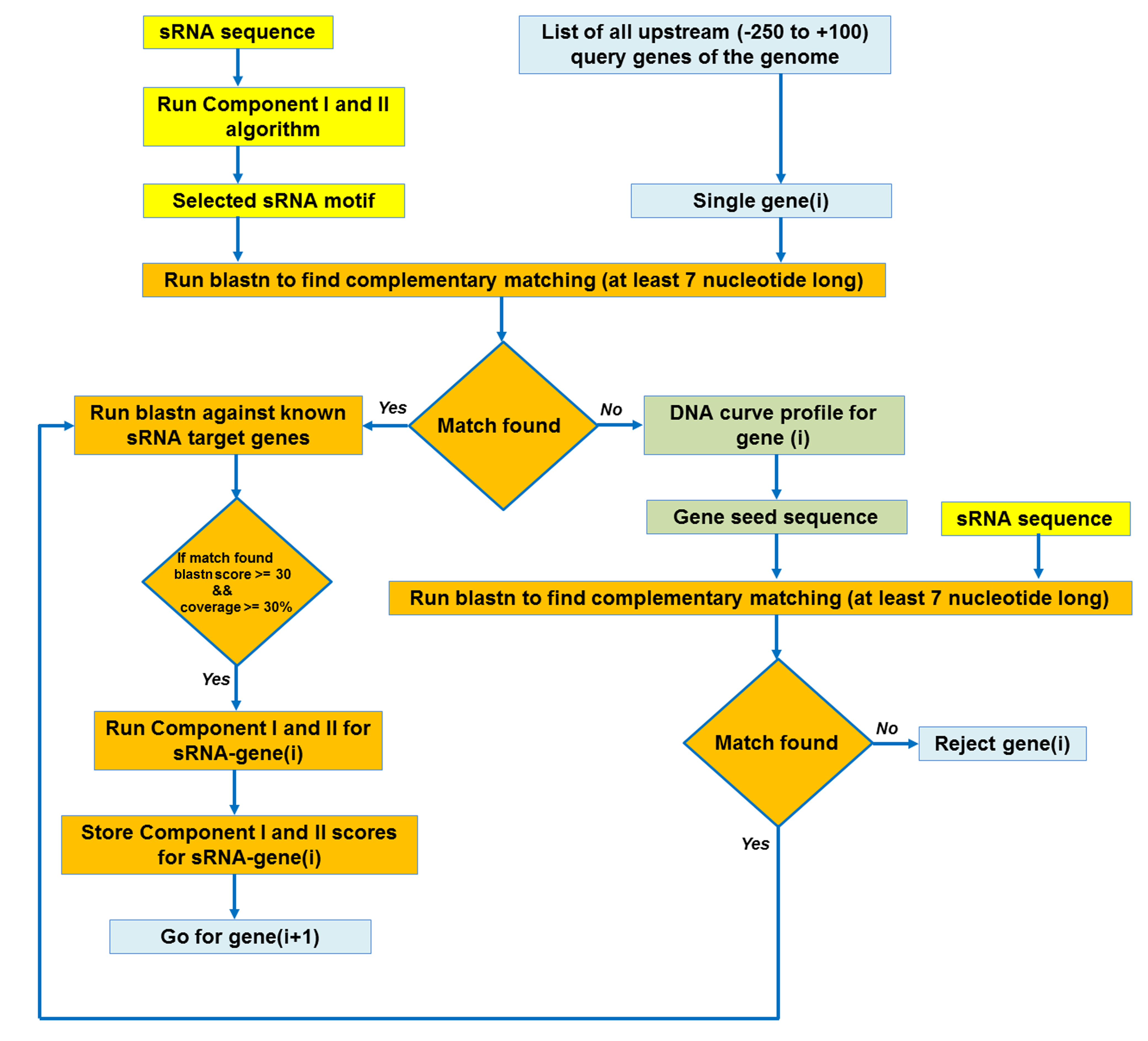

**Figure S2.** Schematic representation of the protocol followed by PresRAT to identify the target genes of sRNA sequences from bacterial chromosome.

**Figure S3**

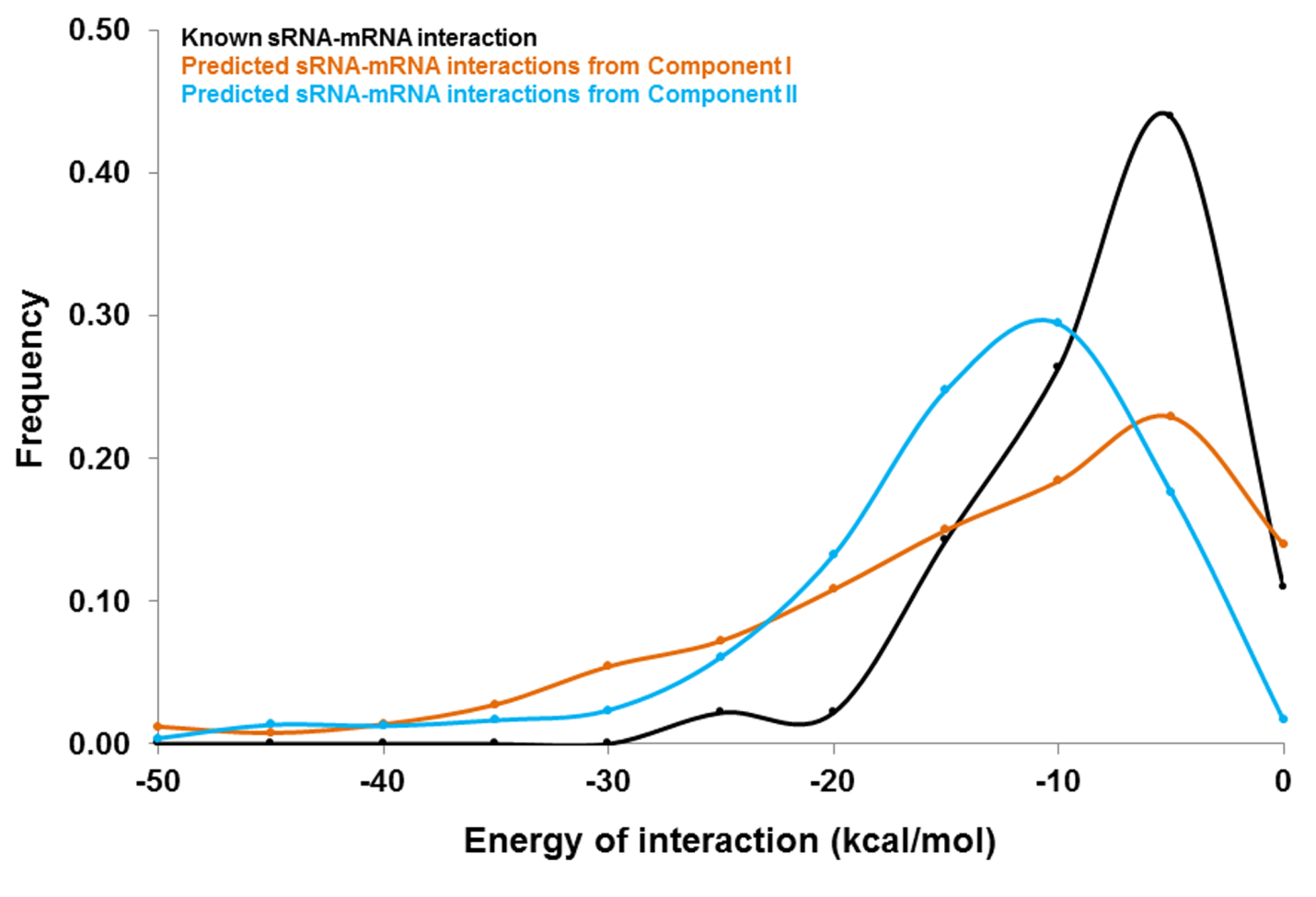

**Figure S3.** Binding energy distribution of known sRNA-mRNA interactions (91 pairs) with predicted sRNA-mRNA interactions (2447 pairs) from both Component I and Component II approaches.

**Figure S4**

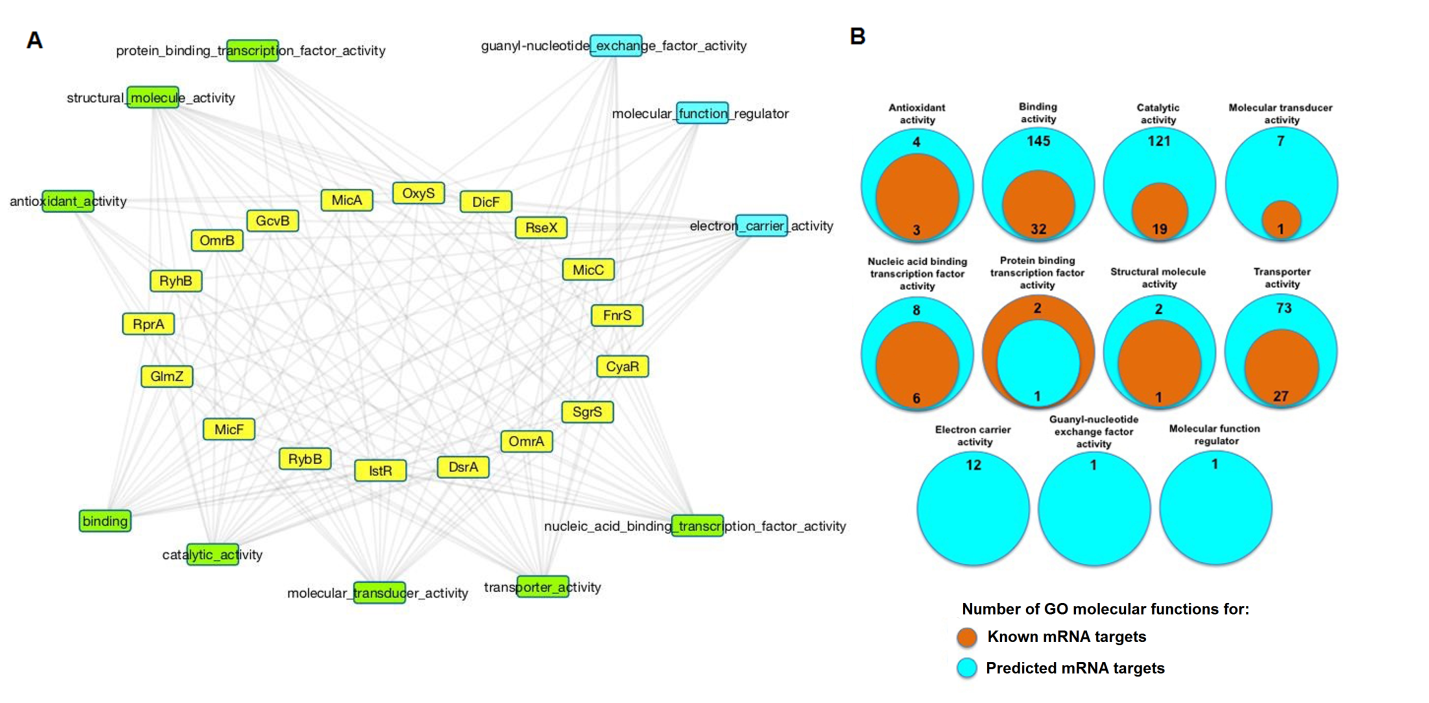

**Figure S4.** Panel A shows the interaction network of 18 *E. coli* sRNA sequences with the molecular functions of their target mRNAs. GO molecular functions were obtained for both the known and predicted mRNA sequences. The yellow nodes represent the sRNA sequences; green nodes represent common GO molecular functions obtained for both known and predicted targets while the cyan nodes represent the exclusive GO molecular functions obtained for the predicted targets. Panel B shows the overlap of GO molecular functions for both known and predicted targets. The orange circles show the number of GO molecular function exclusively obtained for known targets while the cyan circles represent the GO molecular functional information obtained for the predicted targets.

**RNA 3D modelling and molecular dynamic simulation**

To analyse the sRNA and the sRNA-mRNA duplexes in more details, we have integrated a platform to generate the three-dimensional model of RNA from their respective secondary structures using RNA2D3D^18^ program, which is relatively faster in generating the initial 3D structure. Given secondary structure information of a RNA molecule, RNA2D3D^18^ rapidly produces a first-order approximation of the three-dimensional information. Generation of the three-dimensional model is followed by molecular dynamics (MD) based minimization and refinement in presence of water and force-field for a better understanding of the structural characteristics of the sRNA/sRNA-mRNA duplex systems. The MD simulations are performed using GROMACS v4.6.3^19^ program. The AMBER99 force field^20^ is used to describe the sRNA/mRNA systems in presence of SPC water model. Periodic boundary condition is applied to minimize edge effects in the solvated structure. The simulation protocol involved energy minimization of the system using steepest descent followed by conjugate gradient algorithms. To ensure minimum disturbance to the starting structure, the equilibration dynamics of the systems is carried out for 100 picoseconds. Leap-frog algorithm is used to integrate Newton’s equations of motion with an integration time-step of 2 femtoseconds. Twin-range cut-offs of 9 Å and 14 Å is applied to calculate the van der Waals interactions and the neighbour list was updated every five steps. The electrostatics is calculated using Reaction-field method. The final MD simulations are performed for 1000 picoseconds.

***Bacterial small-RNA structural analysis***

An in-depth analysis of sRNAs and their dynamics in presence of water and other proteins is critical to extend our understanding on these functional RNAs. Three-dimensional modelling of sRNA sequences using their existing secondary structure information provides us the opportunity to decipher the properties of these functional RNAs using advanced computational analyses. PresRAT uses existing RNA2D3D^18^ program to build the initial three-dimensional model of sRNA sequences followed by extensive refinements by molecular dynamics simulations as described in the methodology section. Using this automated procedure we have successfully built 50 sRNA models (Table 1). The models were further extensively energy minimized, equilibrated (100 picoseconds) and simulated (1 nanoseconds) in presence of water. The qualities of these refined sRNA 3D models were assessed using WebRASP^21^ program. Negative normalized energy values obtained from WebRASP^21^ (Figure S5A) demonstrated an accurate and stable RNA models. Structural properties such as compactness (radius of gyration), average number of hydrogen bond and entropy of the sRNA structures show a general trend of improvement in values with the growing number of atoms in the sRNA structure (Figure S5B-S5D). The average free energy (∆G) of sRNA structures plotted against its respective sRNA lengths shows (Figure S5E) that all the structures possess negative ∆G values. The ∆G distribution of all the simulated ensemble of sRNA structures (Figure S5F) has a mean free energy of -21 Kcal/mol while the upper and the lower limit of free energy values lie from 0 to -40 Kcal/mol.

**Table 1**

| **sRNA Name** | **Genome** | **Length** |
| --- | --- | --- |
| ChiX | NC_003197 | 80 |
| CyaR | NC_000913 | 87 |
| DicF | NC_000913 | 53 |
| DsrA | NC_000913 | 87 |
| DsrA | NC_003197 | 90 |
| FnrS | NC_000913 | 122 |
| FsrA | NC_000964 | 46 |
| GadY | NC_000913 | 105 |
| InvR | NC_003197 | 93 |
| MicA | NC_000913 | 78 |
| MicA | NC_003197 | 79 |
| MicA | NC_003198 | 76 |
| MicC | NC_003197 | 109 |
| MicF | NC_000913 | 93 |
| MicF | NC_003197 | 93 |
| OmrA | NC_000913 | 88 |
| OmrA | NC_003197 | 87 |
| OmrA | NC_011601 | 88 |
| OmrB | NC_000913 | 82 |
| OmrB | NC_003197 | 85 |
| OmrB | NC_011601 | 76 |
| OxyS | NC_003197 | 121 |
| Qrr1 | NC_002505 | 96 |
| Qrr1 | NC_009783 | 94 |
| Qrr2 | NC_002506 | 108 |
| Qrr2 | NC_009784 | 109 |
| Qrr3 | NC_002506 | 107 |
| Qrr3 | NC_009784 | 109 |
| Qrr4 | NC_002506 | 107 |
| Qrr4 | NC_009784 | 109 |
| RdlD | NC_000913 | 64 |
| RprA | NC_000913 | 106 |
| RprA | NC_003197 | 106 |
| RseX | NC_000913 | 91 |
| RsmY | NC_012660 | 118 |
| RsmZ | NC_002516 | 116 |
| RybB | NC_000913 | 79 |
| RydC | NC_000913 | 64 |
| RydC | NC_003197 | 63 |
| RyhB-1 | NC_003197 | 95 |
| RyhB-2 | NC_003197 | 113 |
| RyhB | NC_000913 | 90 |
| SokB | NC_000913 | 56 |
| SokC | NC_000913 | 55 |
| spf | NC_003197 | 66 |
| SPOT42 | NC_000913 | 109 |
| SPOT42 | NC_011601 | 109 |
| SroB | NC_003197 | 81 |
| SymR | NC_000913 | 77 |
| Yfr1 | NC_005072 | 56 |

**Figure S5**

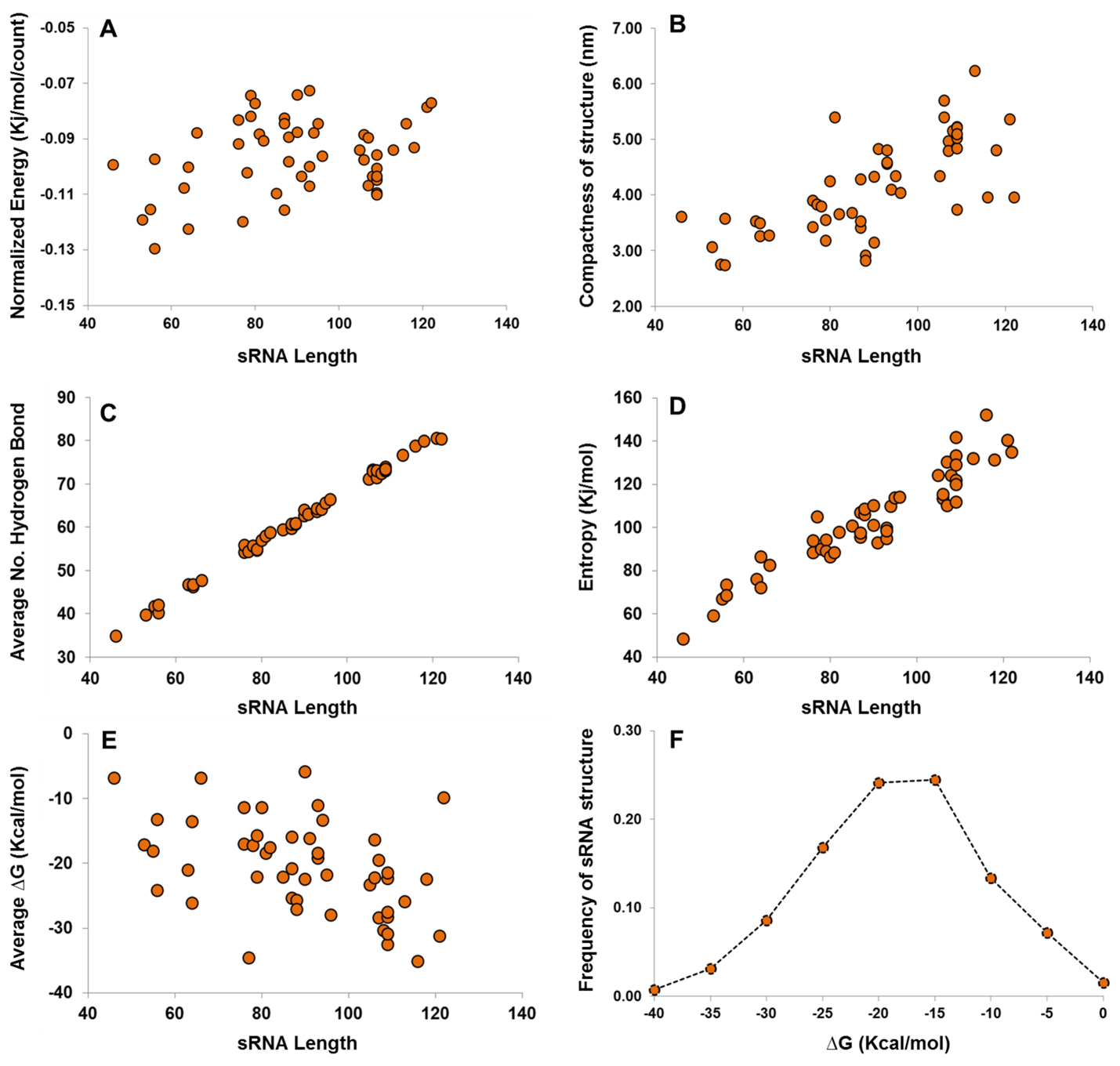

**Figure S5**. The figure depicts the structural properties of 50 refined sRNA models. Panel A shows the normalized energy score calculated using WebRASP^21^ program. The negative potential energy scores obtained by all the refined models describe their good reliability as a whole. Panel B demonstrates the distribution of radius of gyration or the compactness of the 50 sRNA models with respect to its length. Panel C and D illustrate the total number of hydrogen bond and entropy values of the sRNA models with respect to their length. Panel E shows the ΔG (Kcal/mol) values of the sRNA models calculated from their secondary structure information. The distribution suggests that all the 50 structures has average negative free energy values while panel F shows the complete distribution of ΔG values of all the individual conformations computed from the molecular dynamics simulation studies performed on 50 sRNA structures.

***Bacterial sRNA-mRNA duplex structural analysis***

81 experimentally known sRNA-mRNA duplex binding regions were modelled using RNA2D3D^18^ program utilizing their secondary structural information. The models were further energy minimized, equilibrated (100 picoseconds) and simulated (1 nanoseconds) in presence of water. Quality assessment of these models were also done using WebRASP21 platform. Negative values obtained (Figure S6A) from WebRASP21 shows accurate and stable duplex 3D models. Analysis of the ensemble duplex structures reveals that their compactness (radius of gyration), average number of hydrogen bond and entropy increases with the growing number of atoms in the sRNA-mRNA duplex structure (Figure S6B-S6D). The average ∆G values of duplex structures in Figure S6E show a mixed distribution, having both positive and negative values. For most of the ensemble duplex structures a positive ∆G value (mean 8.22 Kcal/mol, Figure S6F) is observed. This observation is curious and indicating that the optimal binding of the sRNA-mRNA region is possibly facilitated by some other factors such as RNA chaperone protein (*e.g.,* Hfq) and/or the other parts of the sRNA, mRNA structure.

**Figure S6**

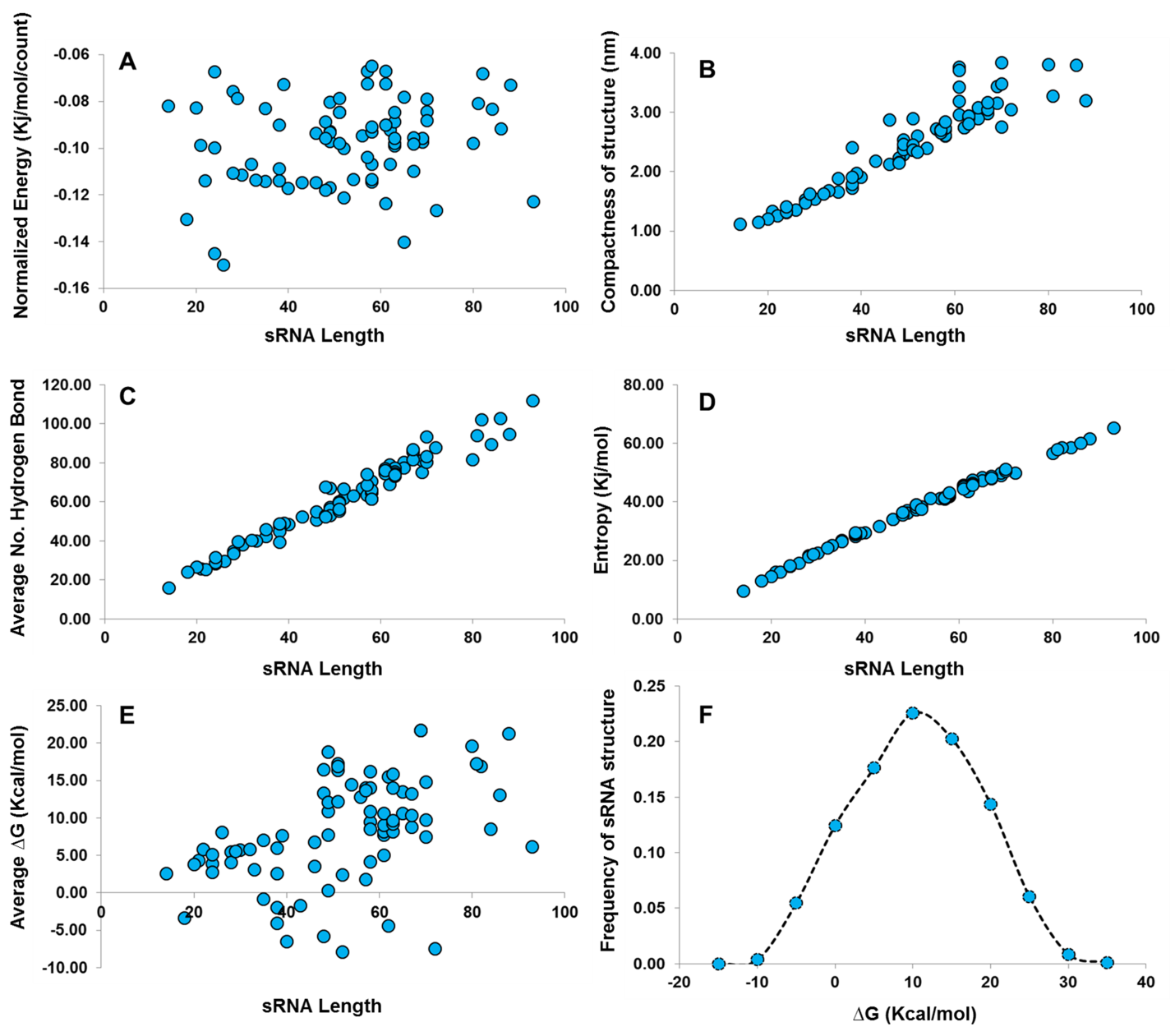

**Figure S6.** Structural properties of 81 refined sRNA-mRNA bound duplex models. Panel A shows the normalized energy score calculated using WebRASP^21^ program. The negative potential energy scores obtained by all the refined models describe their good reliability as a whole. Panel B demonstrates the distribution of radius of gyration (compactness) of the 81 sRNA-mRNA bound duplex models with respect to bound sRNA length. Panel C and D illustrate the total number of hydrogen bond and entropy values of the sRNA-mRNA bound duplex models with respect to their length. Panel E shows the ΔG (Kcal/mol) values of the sRNA-mRNA bound duplex models calculated from their secondary structure information. Panel F shows the complete distribution of ΔG values of all the individual conformations computed from the molecular dynamics simulation studies performed on 81 sRNA-mRNA bound duplex structures.

**Figure S7**

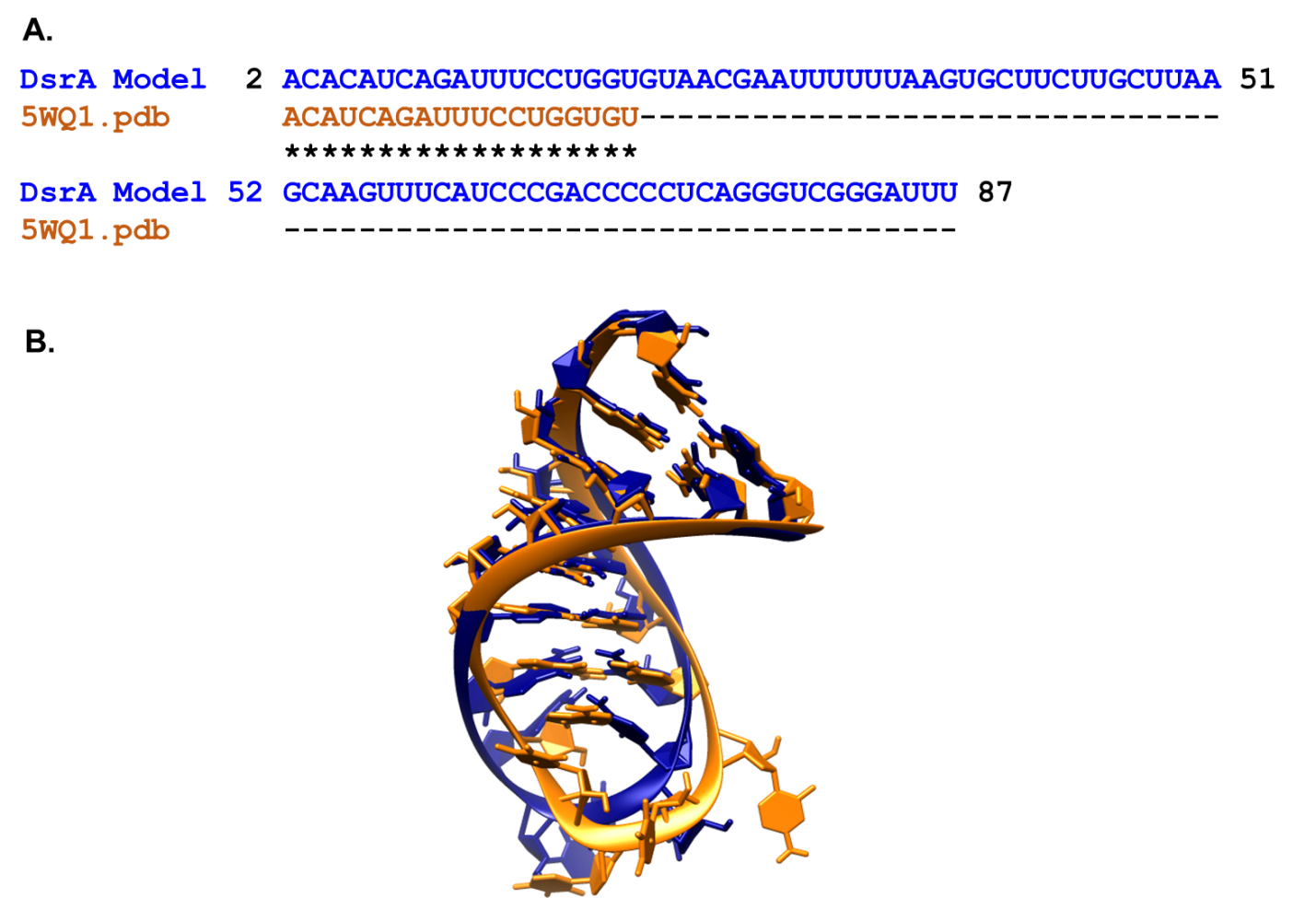

**Figure S7**. Comparison of sequence and structural similarity between experimental NMR structure and computationally derived 3D model of DsrA. Panel A show the sequence alignment between the NMR fraction of DsrA (golden colour) and the PresRAT derived 3D model of DsrA (blue colour) whereas panel B shows the structural superimposition between the two structures.
